# High Resolution Comparison of Cancer-Related Developmental Processes Using Trajectory Alignment

**DOI:** 10.1101/469601

**Authors:** Ayelet Alpert, Elina Starosvetsky, Michal Hayun, Yishai Ofran, Shai S. Shen-Orr

**Author notes:** These authors jointly directed this work (Yishai Ofran, Shai S. Shen-Orr). Correspondence should be sent to or.

## Abstract

Abnormal differentiation is a key feature of cancer, yet currently there is no framework that enables a comparative analysis of differentiation processes across patients while preserving their individual-level resolution. Here, we present devMap, an algorithm that uses high-dimensional trajectory alignment to anchor cancer-related developmental processes to a common backbone process, thus allowing for their systematic comparison. We applied devMap to bone marrow samples from healthy individuals and AML patients profiled by single-cell mass-cytometry at cancer diagnosis and following treatment. devMap standardization enabled us to infer the developmental status of the AML samples and characterize its evolution following treatment and in relapse. Application of devMap on an external dataset of AML bone marrow samples revealed conserved patterns of developmental signaling responses in AML that were obscured by traditional methodologies for developmental inference.

## Introduction

Cellular stoichiometry is highly conserved in the healthy tissue due to tightly regulated cellular differentiation processes. However, in cancer, mutations in key transcription factors lead to a partial removal of these regulations, resulting with unbalanced cellular frequencies commonly observed as an expansion of a relatively homogeneous cell population, a clone of cells from a certain developmental stage. In clinics, this developmental stage is regularly assessed via morphologic or phenotypic similarity between the cells of the major clone in the tumor and the healthy cells, information that is subsequently used for cancer classification. However, growing evidence suggests that despite being the dominant population in the tumor, the expanded clone is only an indirect consequence of cancer pathogenesis, whereas the cellular origin of cancer is attributed to cells from very early developmental stages. This highlights the necessity in characterization of the entire differentiation process occurring in the cancerous lineage.

Normal differentiation and developmental processes have been recently studied in the context of developmental trajectories, a computational tool that uses snapshot high-dimensional data collected on single cells to order them along a continuum capturing their developmental stage. By using inter-cellular similarity as a metric, developmental trajectories enable characterization of markers expression dynamics along the process at unprecedentedly high temporal resolution. The conservation of developmental processes between healthy individuals allowed for averaging of multiple samples taken from different individuals to assemble one common trajectory. However, in cancer, the high variability existing between patients with respect to the developmental alterations hinders such an averaging, resulting with patient-specific trajectories. While being useful for understanding the unique process occurring within a certain patient, lack of tools for comparison between these trajectories prevents a global and systematic characterization of cancer-related developmental alterations.

While these principles apply to numerous cancers, they are highly relevant in acute myeloid leukemia (AML), the most common acute leukemia in adulthood^2^, in which myeloid progenitor cells spanning various developmental stages (blasts) are accumulated in the bone marrow^1^. Traditional classification systems use the blasts’ morphology to classify AML cases, with the resulting subtypes linked to specific mutations^4^ and thereby are closely related to clinical outcome^5^. However, by focusing on the dominant clone, this approach ignores other aspects of the entire differentiation process occurring in the cancer that may be associated with its cellular mechanism and thus potentially include clinically relevant information.

Here, we present the devMap, an algorithm that robustly aligns cancer-specific trajectories to the healthy backbone trajectory, allowing for their comparison across patients and against the healthy counterpart. To showcase the utility of devMap, we applied it on a dataset of healthy and AML bone marrow samples profiled by mass cytometry for myeloid-maturation related markers. We assembled the AML trajectories in a patient-specific manner and aligned them using devMap to the conserved monocytic developmental trajectory obtained by averaging healthy samples. The standardization achieved by this approach was leveraged to compare the developmental stages of AML samples that was associated with clinical staging. Mapping of an external dataset onto the healthy trajectory revealed a conservation of maturation dependent signaling responses in AML that was obscured when using traditional methodologies for developmental inference.

## Results

### Averaging of cancer samples for trajectory assembly results in a substantial loss of the patient-level resolution as compared to healthy

The high conservation of developmental processes within healthy individuals yielded the traditional averaging approach, where researchers generate one trajectory after pooling of multiple healthy samples to characterize the common process. However, in cancer, unique genetic alterations occurring within a patient often yield changes in markers expression dynamics along these developmental processes, supporting a patient-specific trajectory model. While trajectory assembly individually for each patient may shed light on the alterations occurring within the patient, such an approach hinders a systematic comparison between developmental processes occurring within different patients.

To illustrate this problem, we used acute myeloid leukemia (AML) as a model of cancer that supports a hierarchical development, where normal differentiation may be disrupted at multiple stages. For characterization of developmental alterations in AML as compared to the healthy, we used CyTOF to profile at a single cell resolution human bone marrow samples taken from nine healthy individuals and 17 AML patients that were sampled longitudinally at the time of diagnosis and 14 days following initiation of anti-cancer treatment. For five patients, we profiled an additional bone marrow sample from the time of relapse diagnosis (Fig. 1A, Methods, **Supp. Tables 1,2**).

**Figure 1.**
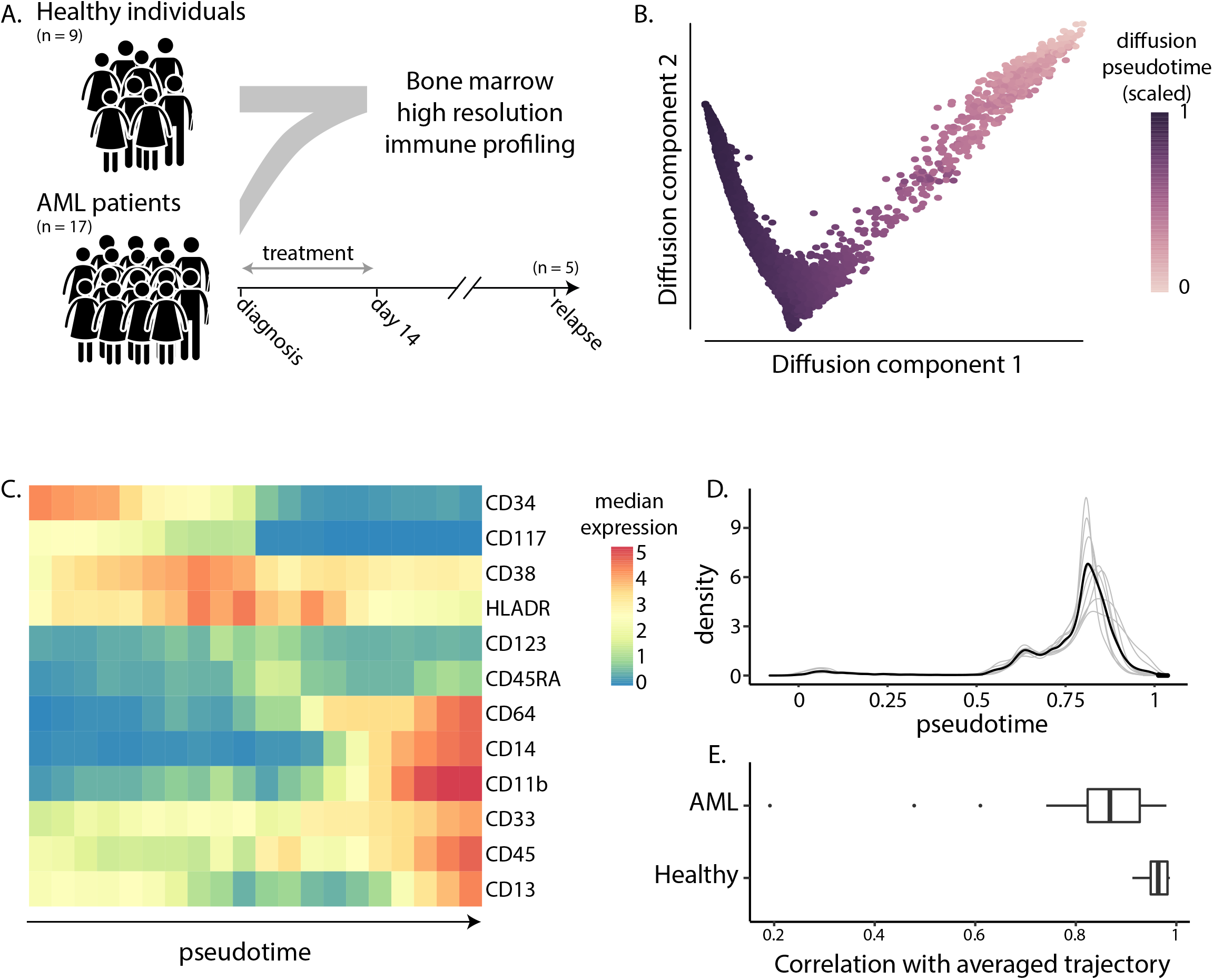
High resolution profiling of bone marrow samples from healthy individuals reveals a conserved trajectory of monocytic development. (A) High resolution profiling of human bone marrow samples of 9 healthy individuals and 17 AML patients sampled longitudinally at the me of diagnosis and 14 days after treatment. For five patients, an additional bone marrow relapse sample was profiled. (B) Diffusion maps 2-dimensional projection of single cells using myeloid markers. Single cells are colored based on the assigned diffusion pseudotime scores. (C) Median markers expression by cells grouped by their positions along the pseudotime axis. (D) Density distribu ons of cells along the pseudotime axis of each healthy individual (grey lines) and averaged across all healthy individuals (black line). (E) Correlations between pseudotime values calculated using either pooled or individual samples of the healthy (lower panel) and AML (upper panel) cohorts (t test P = 1.7 ∙10^−3^).

We first aimed to determine the conservation of developmental processes occurring within healthy individuals. For this, we applied the diffusion pseudotime algorithm^20^, an algorithm regularly used to order cells along developmental trajectories, on all the 9 healthy bone marrow samples to reconstruct the healthy monocytic differentiation from the early stem cells to mature monocytes (Fig. 1B, Methods). Markers expression dynamics along the trajectory matched their known dynamics during monocytic differentiation, with downregulation of progenitors-associated markers such as CD117 and CD34, and upregulation of mature-monocytes associated markers such as CD14 and CD11b (Fig. 1C). We thus concluded that the assembled averaged trajectory reflects normal monocytic maturation.

We next sought to study the conservation of this process among our healthy cohort by comparison of the individual-level cellular density along the trajectory. Despite populating the entire range of the trajectory, all healthy individuals exhibited two density peaks corresponding to stable states: a high peak of mature monocytes and a smaller one of immature blasts, suggesting an overall conservation of the stochiometric relationships between monocytic cell populations in the healthy bone marrow (Fig. 1D).

Next, to determine the conservation in markers expression dynamics throughout the process across individuals, we assembled a trajectory for each healthy individual separately and compared it to the averaged one. The correlations between the resulting pseudotime values obtained by both approaches were high (0.96±0.025), justifying the averaging of multiple samples for achieving a robust characterization of the normal developmental process (Fig. 1E). However, when we applied a similar approach to the diagnosis and relapse cancer samples in the AML cohort, we observed significantly lower correlations between the pseudotime values obtained using these approaches (t test P = 1.7∙10^−3^, 0.81±0.183, Fig. 1E). This indicates that while providing a comparative framework of developmental processes occurring in different patients, an assembly of an averaged trajectory through pooling of multiple cancer samples results with a substantial loss of the patient-specific information.

### The devMap algorithm aligns cancer-related developmental processes onto the healthy counterpart by trajectory alignment

To resolve this, we developed the devMap algorithm, a developmental mapper that enables a quantitative comparison of developmental processes assembled individually, thus preserving the individual-level resolution of the process. To accomplish this, devMap uses single-cell trajectory alignment in two sequential stages to achieve a common scaling of the patients-derived trajectories, allowing for their comparison. Specifically, devMap takes as input cancer trajectories assembled for each patient individually and an averaged healthy trajectory that will serve as a reference backbone. For each patient, devMap first identifies the set of markers exhibiting conserved expression dynamics along the trajectory as compared to the healthy backbone by minimizing the alignment cost of these two trajectories. devMap then uses this patient-specific set of conserved markers to re-align the patient-derived trajectory to the healthy backbone and leverages this alignment to scale the patient-derived pseudotime axis based on the reference trajectory. The resulting common scaling of the patients’ derived trajectories enables their systematic comparison across patients in a downstream analysis (Fig. 2, Methods).

**Figure 2.**
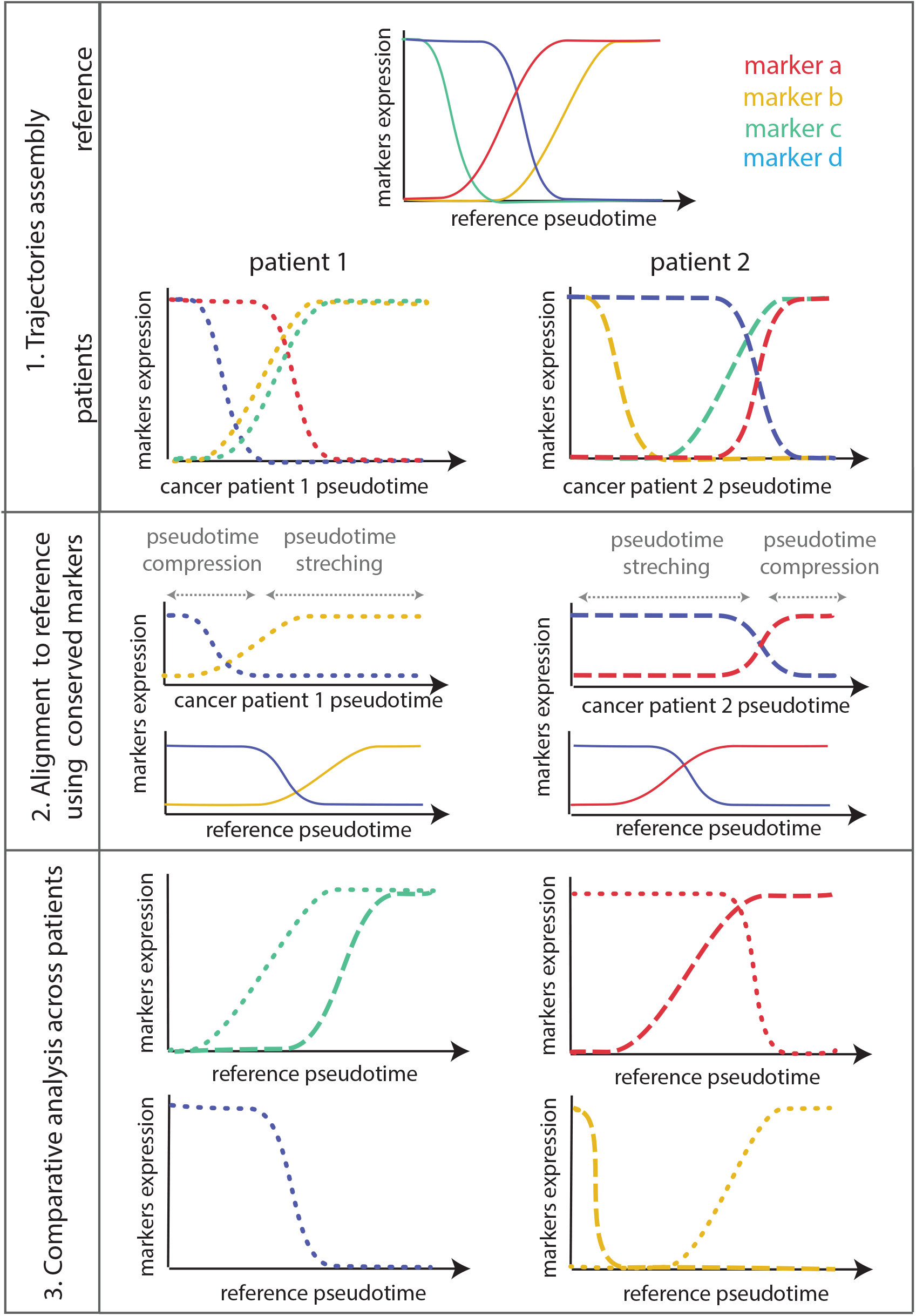
Overview of devMap algorithm. devMap takes as input pseudotime trajectories assembled from the healthy reference and cancer patients (upper panel). devMap then identifies the markers with conserved expression dynamics along the cancer-derived trajectories as compared to the healthy and uses them for high-dimensional alignment of these trajectories (middle panel). The resulting alignment is used to scale the pseudotime axes of the cancer-derived trajectories based on the healthy reference (lower panel), allowing for a marker-wise comparative analysis.

To validate devMap performance, we applied it on single-cell trajectories of AML patients and healthy individuals using either real or synthetic data, where the ground-truth is known (Fig. 3A). The first synthetic AML dataset was generated by repeated subsampling of healthy bone marrow monocytes with a sampling bias towards different locations along the healthy trajectory, to produce altered cellular density distributions along the trajectory. The biased sampling influenced the trajectory’s curvature as reflected by deviations of the assigned pseudotime values of cells in the synthetic data from their original values. These deviations were significantly diminished following devMap application, establishing the ability of devMap to overcome cellular density abnormalities that often characterize cancer (Fig. 3B, Methods).

**Figure 3.**
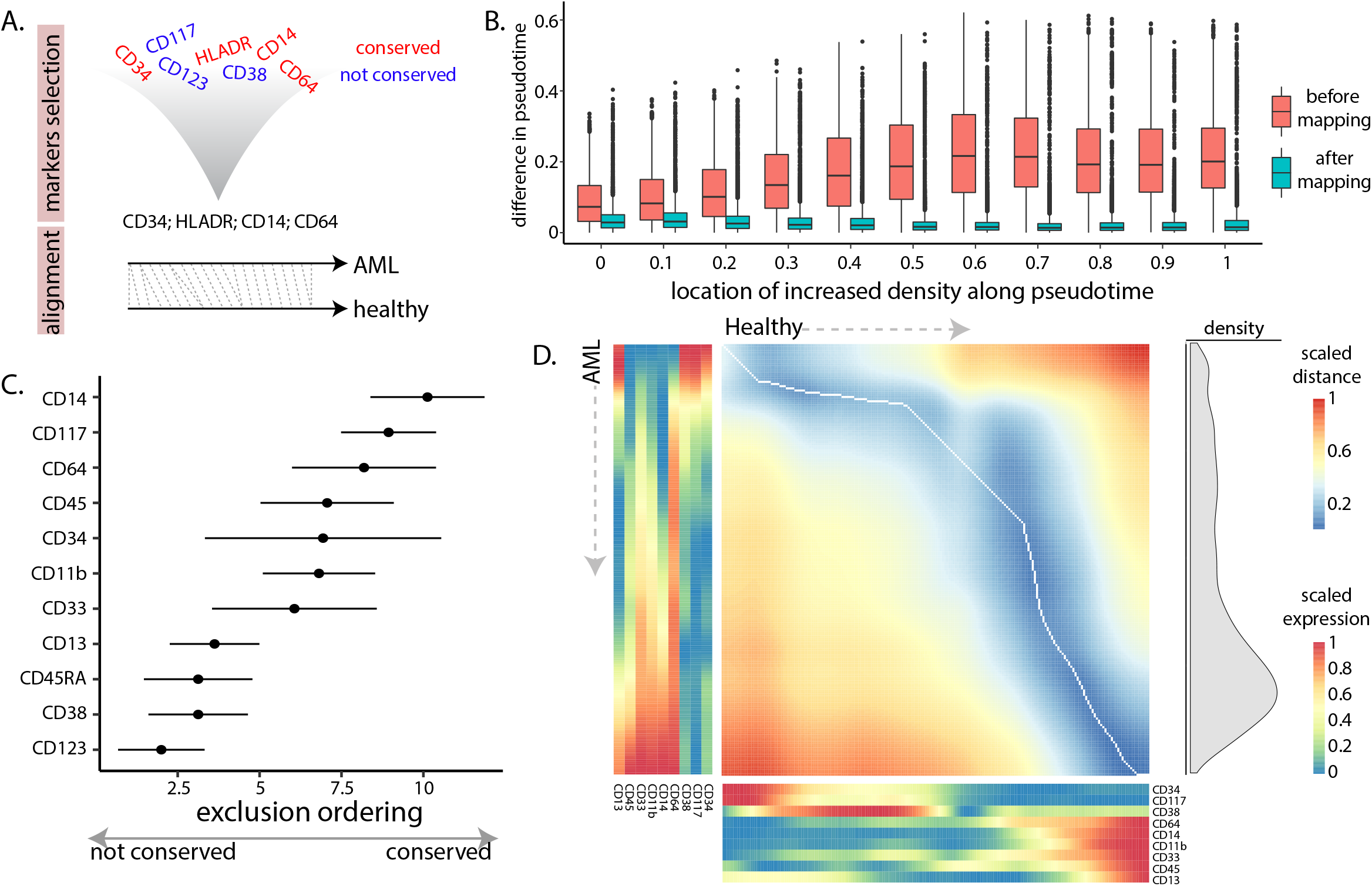
Application of devMap on synthe c and real datasets of AML for performance demonstration. (A) Following myeloid markers selection relying on the conserva on of expression dynamics along the AML trajectory as compared to the healthy trajectory, the two trajectories are aligned. (B) Increased cellular density in different locations along the healthy trajectory changes the assigned pseudotime values (pink boxes), that are restored back to their original values upon devMap application (blue boxes). (C) Exclusion ordering of markers across patients as determined by their dynamics conserva on along the AML trajectory compared to the healthy trajectory. Dots correspond to median whereas error bars correspond to standard deviations, both were calculated across patients. (D) Dissimilarity matrix and global alignment path between the healthy (left to right) and AML (top to bo om) trajectories using the selected set of markers. Interpolated and scaled expression dynamics of those markers selected for the alignment along the healthy and AML trajectories appear to the bottom and left, accordingly. Density of cells along the AML trajectory appears to the right.

Next, to validate devMap’s ability to identify abnormalities in markers expression dynamics to extract a relevant set of markers for the alignment, we generated for each marker an additional synthetic dataset in which the expression dynamics of the marker was reversed. For all the markers, devMap correctly identified the marker with the altered expression dynamics as the one whose exclusion is the most beneficial for the alignment quality. We thus concluded that by identification of a set of markers with conserved expression dynamics, devMap’s alignment can overcome cellular density abnormalities and standardize the pseudotime axes of cancer samples.

We thus applied devMap on our dataset to align AML patients’ trajectories to the averaged healthy trajectory. Summarizing the markers’ exclusion ordering across patients revealed the relative tendency of the different markers to preserve their expression dynamics in our cohort of AML patients (Fig. 3C). For four patients, the selected set of markers that could achieve a high-quality alignment included less than five markers, indicating that markers expression dynamics along the AML trajectories of these patients were extremely abnormal, and thus we excluded them from further analysis. The remaining patients varied by the number of markers used for alignment, with 9 as the median number of markers with conserved expression. devMap’s markers selection step improved the alignment quality, allowing for a meaningful scaling of the patient-derived pseudotime curvature based on the corresponding healthy trajectory (Fig. 3D).

### devMap application enables inference of cellular maturation state of cancer samples to reveal AML evolution under treatment and in relapse

devMap alignment of cancer-related trajectories to the common healthy reference can be used to standardize the pseudotime curvatures across patients, allowing for their systematic comparison. We sought to leverage this standardization to generate a metric estimating the overall maturation state of the cells in a sample as reflected by the cellular density distribution. Specifically, we defined the maturation index of a sample as the area under the cumulative density distribution curve, with high values correspond to an enrichment of immature cells whereas low values correspond to an enrichment of mature cells (Fig. 4A).

**Figure 4.**
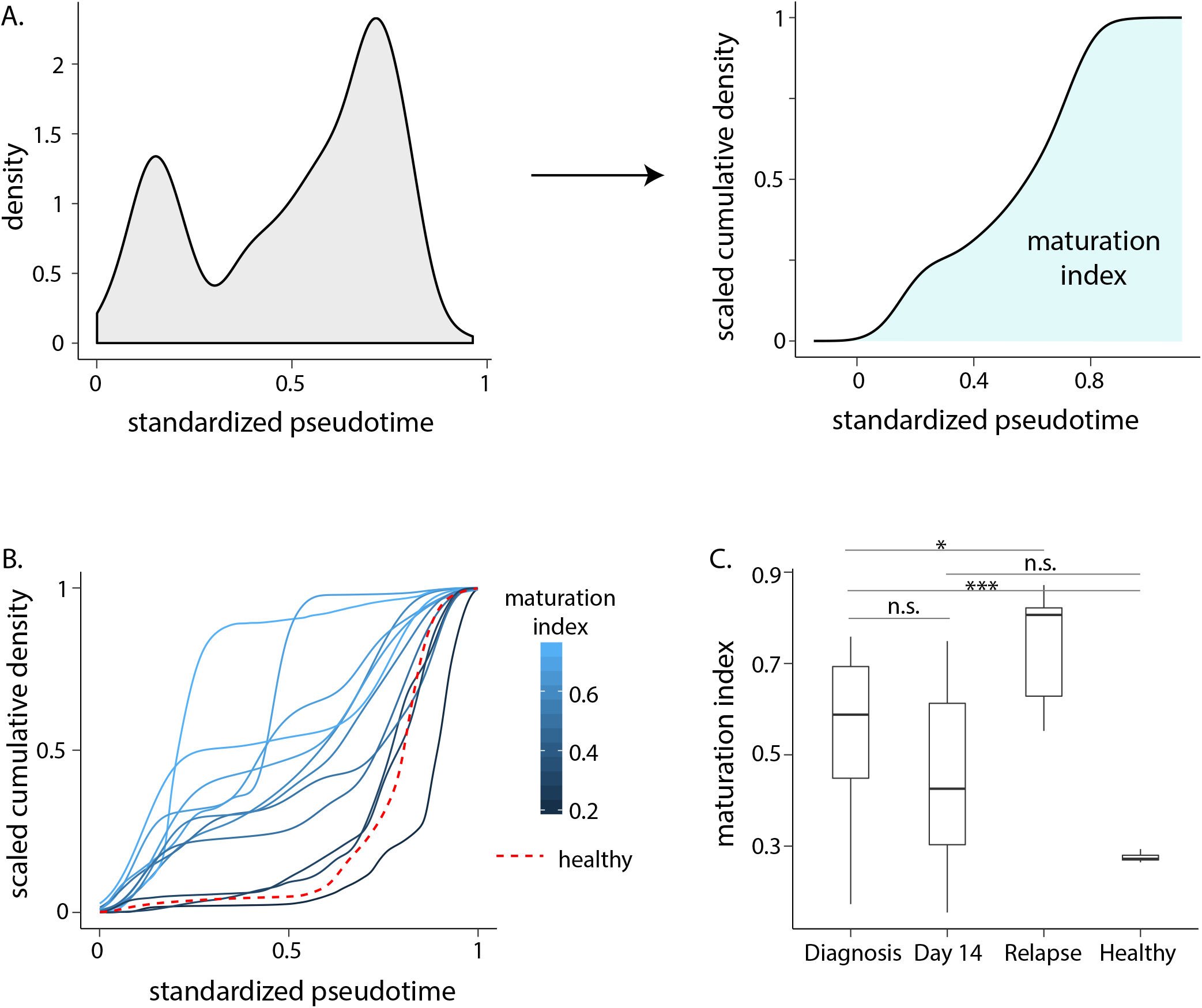
Standardization of pseudotime axis enables inference of the overall cellular matura on degree of a sample. (A) Density distribution of cells along the standardized pseudotime axis (left) can be transformed into a scaled cumulative distribution (right). The area under the scaled cumulative distribution line represents the overall density along the scaled trajectory, namely: a maturation index. (B) Scaled cumulative density distributions of different patients along the standardized pseudotime axis colored by the calculated maturation index. The mean corresponding curve of healthy individuals appears as a dashed red line. (C) Maturation indices calculated across healthy individuals (right-most bar) or AML patients at different stages of their disease. *,*** p value < 0.05, 0.001 by t test; n.s.: not significant.

The cumulative cellular density distributions varied among the AML patients in our cohort with respect to both the quantity and the locations of cellular density peaks, corresponding to the number of clones and their relative maturation, respectively (Fig. 4B). The maturation indices of AML patients at diagnosis were significantly higher than those calculated on healthy individuals (p value = 5∙10^−4^), reflecting the incomplete maturation commonly observed in cancer. However, the maturation indices of those samples collected 14 days after treatment were lower than those calculated on the diagnosis samples, yet higher compared to the healthy, suggesting that in our cohort a complete response was not always achieved following 14 days of treatment (Fig. 4C). To characterize the dynamics of cellular maturation in AML, we used those five patients for which additional bone marrow samples were collected at the time of AML relapse. The calculated AML maturation indices were significantly higher in the relapse samples as compared to the initial diagnosis, indicating that in agreement with a previous report^21^, AML recurrence tends to involve earlier maturation stages than the initial diagnosis (Fig. 4C).

### Application of devMap on an external dataset reveals a novel conservation of developmental signaling dynamics in AML that was obscured under the traditional methodology

In addition to the altered cellular differentiation, abnormal signaling responses have been previously reported to occur in AML^9,10^. To characterize them in a developmental context, we utilized the dataset generated by Levin et al^9^ that profiled a cohort of five healthy individuals and 16 AML patients using CyTOF by measuring the abundance of extracellular myeloid epitopes along with intracellular signaling molecules at baseline and following 16 perturbations.

To test devMap ability to overcome the technical differences between the datasets, we applied it using the common extracellular myeloid markers measured in both datasets to scale the pseudotime curvature of the individual healthy samples from the external dataset to our healthy reference (Methods, Supp. Fig. 3). For each individual, we calculated the maturation index either before and following pseudotime standardization by devMap and correlated it with the corresponding frequency of immature cells expressing CD34 in the sample, as obtained by manual gating. By anchoring the individual-level healthy trajectories to our common reference, devMap application unraveled the underlying correspondence between the maturation index and the frequency of CD34^+^ cells (Fig. 5A, Pearson correlations: 0.34, 0.99, p values: 0.54, 0.001, calculated either without or with mapping, respectively). We thus concluded that by overcoming the technical variability between the datasets, devMap can be used to characterize signaling responses dynamics throughout monocytic development in healthy individuals and AML patients.

**Figure 5.**
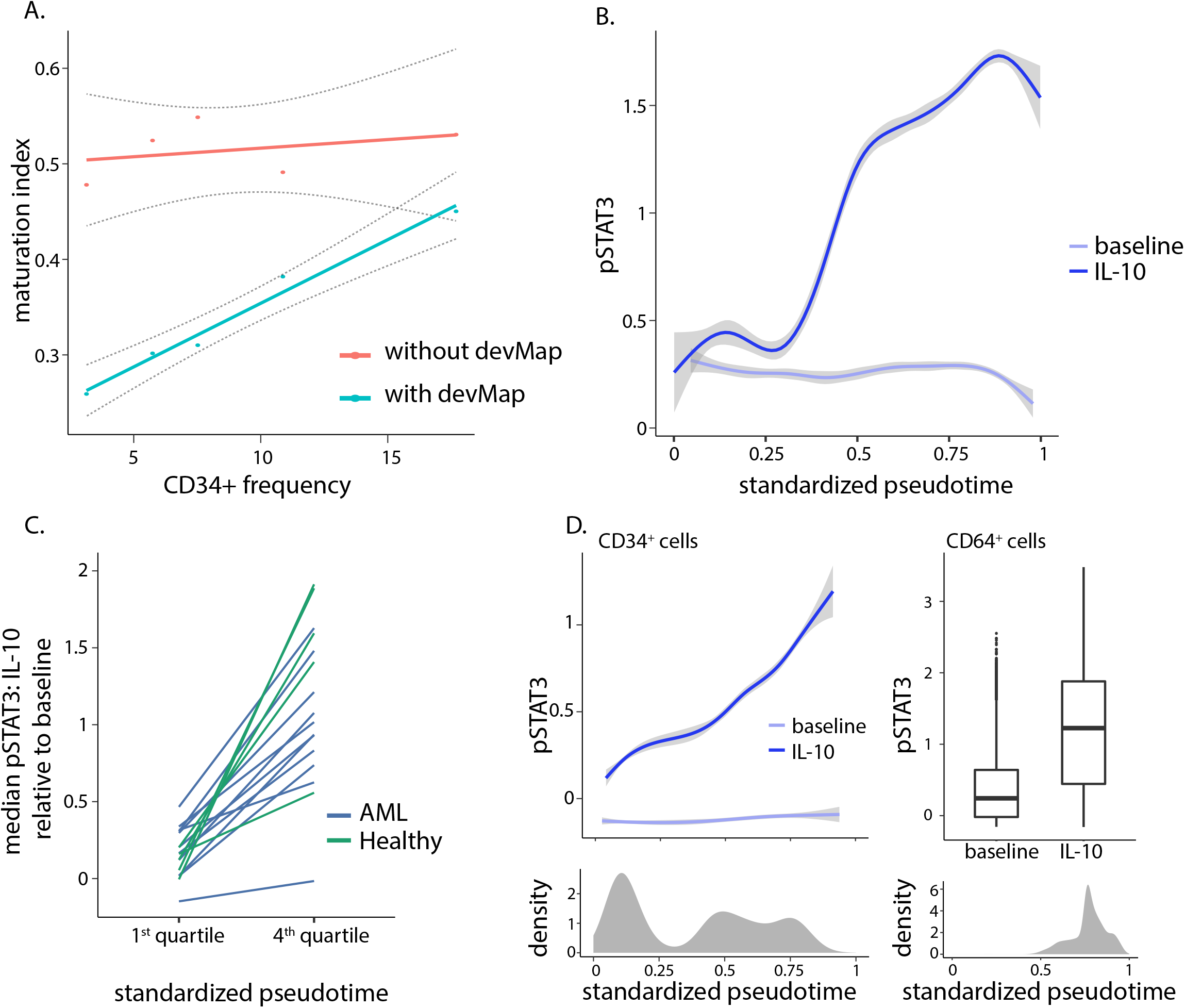
Cellular mapping onto the healthy trajectory overcomes technical differences between datasets generated by different labs and is essen al for correct inference of cellular maturation state. (A) Frequencies of the manually gated CD34^+^ cell popula on measured in healthy individuals measured by Levin et al against the maturation indices calculated either without (pink) or with (blue) devMap application. (B) Dynamics of phosphorylated STAT3 expression along the standardized pseudotime of one healthy individual at baseline (light blue) and following IL-10 s mula on (blue). (C) Median expression of phosphorylated STAT3 under IL-10 stimulation relative to baseline as calculated using the cells assigned to the first and forth quartiles of the standardized pseudotime in healthy individuals (green) and AML patients (blue). (D) Top left: Gated CD34^+^ cells exhibit variable STAT3 phosphoryla on responses to IL-10 dictated by their location along the pseudotime axis. Bottom left: The gated CD34^+^ population are broadly distributed along the standardized pseudotime axis. Top right: Gated CD64^+^ cells exhibit a conserved STAT3 phosphoryla on responses to IL-10. Bo om right: The gated CD64^+^ popula on are narrowly distributed along the standardized pseudotime axis.

Four signaling responses were originally identified by Levine et al^9^ as correlated with myeloid maturation in healthy individuals. Specifically, we focused on IL-10 induced STAT3 phosphorylation response that was shown to be normally stronger in differentiated as compared to immature myeloid cells, a correlation that was not consistently preserved in AML patients. To characterize this association in the context of the healthy development, we smoothed the expression of phosphorylated STAT3 (pSTAT3) along the standardized trajectory at baseline and following IL-10 stimulation. In agreement with the original finding, the relative expression of pSTAT3 increased with the pseudotime axis in all healthy individuals, suggesting that devMap can be used to identify maturation-dependent signaling responses (Fig. 5B, **Supp. Fig. 4**).

Next, to explore the dynamics of maturation-dependent signaling in AML, we used devMap to standardize the AML-dependent trajectories based on our healthy reference, each time selecting a marker-set with conserved expression dynamics, resulting with 11 AML samples with reliable mappings (Methods). We observed that IL-10 induced STAT3 phosphorylation was significantly increased in the terminal parts of the standardized trajectories as compared to their beginnings, suggesting a conservation of the correlation of this signaling response with differentiation in AML (Fig. 5C).

devMap enables inference of the developmental state of single cells using the rich high dimensional data, yielding an accurate and robust assessment. To showcase this added value, we compared our approach to the one taken by the original publication that showed an altered expression of CD34 in those cell clusters exhibiting signaling patterns that normally correlate with a mature phenotype. Such an approach solely relies on the expression of one marker as a representative of the whole extracellular expression profile. We identified one patient for whom the stemness marker CD34 was identified by devMap as having altered expression dynamics as compared to the healthy trajectory, and thus was excluded from the alignment. Indeed, in this patient, cells positive for CD34 were mapped to almost the entire range of the pseudotime axis, including regions that correspond to mature cells (Fig. 5D, **bottom left**). Despite expressing CD34, this cell population exhibited signaling dynamics that matched the normal dynamics along the trajectory, where cells that were mapped to less mature parts of the trajectory exhibited weaker responses as compared to those that were mapped to the mature parts (Fig. 5D, **top left**). Conversely, cells positive to the maturation marker CD64, a marker that was included in the alignment process, showed the expected distribution along the trajectory along with a strong IL-10 induced STAT3 phosphorylation (Fig. 5D, **right**). These results indicate that using CD34 as a single representative to cellular maturation state may be misleading in AML, while exploiting the rich high-dimensional information by devMap is essential for an accurate derivation of cellular maturation state.

## Discussion

Unlike healthy processes, cancer related developmental processes exhibit a large inter-patient variability, hindering a combinatorial analysis of multiple samples concomitantly and highlighting the need in a personalized approach. While achieving the high intra-individual resolution, this approach lacks downstream analytical tools that allow for a comparative analysis of developmental processes across patients.

Here we developed devMap, an algorithm that aligns cancer-related developmental processes to one common healthy reference, thus allowing for their standardized comparison. For this, devMap first identifies for each patient only those markers with conserved expression dynamics as compared to the healthy and uses them for a high-dimensional alignment. By relying on a large set of markers for alignment, devMap maps the cancer-related and healthy trajectory accurately and robustly. The resulting alignment can be used both for deriving the approximated developmental stage of single cancer cells, and for comparison of the dynamic behavior of different features, such as signaling responses, along the trajectories of different patients.

We used single-cell mass cytometry data of AML patients to demonstrate the resolution loss resulting from using multiple samples to derive one averaged trajectory in cancer. devMap’s standardization allowed for a comparative analysis of cellular density distributions along the AML-related trajectories, revealing insights regarding AML evolution following treatment and in relapse. Alignment of an external AML single-cell data to the reference trajectory, highlighted the previously obscured conservation of the dynamics of developmental signaling responses in AML, that was revealed due to the high-dimensional alignment approach and markers selection step.

devMap relies on some fundamental features of the cancer sample, namely: the existence of an inferable trajectory within the sample along which a large-enough set of markers exhibit conserved expression dynamics as compared to the reference backbone. Thus, for the minority of cases where the aberrant structure of the cancer sample prevents an accurate assembly of a trajectory or where markers expression dynamics are extremely abnormal, an accurate mapping cannot be derived. In the latter case, the algorithm will suggest that relying on a single or a group of markers to determine single-cell maturation state may be misleading. However, we note that increasing the breadth of the markers measured per cell by more advanced technologies or application of a preliminary step in which the markers are chosen a priory based on their conservation with the healthy might significantly help in reducing these cases.

By aligning samples profiled using multiple methodologies onto the reference backbone, one can enrich the characterization the developmental processes occurring within the cancer sample, overcoming technological limitations with respect to the number of measured features. This enrichment can be used for high dimensional comparison of developmental processes occurring within groups of patients classified based on clinical features, thus may suggest potential molecular mechanisms that can account for the observed clinical differences.

In addition to its impact on basic cancer research, devMap can be widely used in clinics to reveal associations between developmental features and clinical consequences, thus suggesting potential therapeutic strategies. Guided by devMap’s mapping, cells can be meaningfully clustered by their developmental stage, as derived from the healthy reference. Studying different molecular features on these cell clusters can be used to derive clinical associations [Good et al].

In the recent years, high-throughput profiling of cancer samples by single-cell technologies such as single-cell RNA sequencing and mass cytometry continues to grow, allowing for a high-resolution characterization of developmental processes that are frequently abnormal in cancer. Comparative framework of these is an essential tool for patients’ stratification that can potentially shed light on molecular and clinical associations, placing devMap as a promising tool for cancer research.

## Acknowledgements

This work was supported in part by grants from the Israeli Science Foundation (grant 1365/12) and Rappaport Institute award to SSO.

## Authors’ contributions

AA, YO, ES and MH and SSO designed the experiment, AA and SSO developed the algorithm, AA, ES, SSO performed analysis of the data, AA, ES, MH performed experiments, MH, YO collected samples, AA and SSO wrote the manuscript, YO and SSO oversaw the research.

## Methods

### Ethics approval

This study was approved by the Helsinki committee affiliated to Rambam Medical Center, Haifa, Israel (IRB number: 0076-15-RMB). All participants gave a written informed consent.

### CyTOF profiling of healthy and AML bone marrow samples

*Samples collection*. Bone marrow aspirates were collected from healthy and AML patients presenting at the Hematology Department in Rambam Medical Health Care Campus. Mononuclear cells were separated by centrifugation over a layer of LymphoprepTM (Axis-Shield PoC AS, Oslo, Norway) and then stored in freezing medium [fetal bovine serum (FBS) with 10% DMSO] in liquid nitrogen. *CyTOF profiling*. Primary conjugates of mass cytometry antibodies were prepared using the MaxPAR antibody conjugation kit (Fluidigm Inc.) according to the manufacturer protocol and optimal concentration was determined by titration. Bone marrow derived cells were thawed in warm media (10% FCS, 1% PSG) and stained with Rhodium intercalator (Fluidigm Inc., 1:2,000 concentration) to discriminate between dead and live cells. The cells were washed with cell staining media (CSM, PBS + 0.5% BSA) and stained with the antibodies cocktail (antibodies details appear in **Supp. Table 4**) in 100ul at room temperature, followed by another wash, fixation (paraformaldehyde 1.6%, in 1ml, RT), and Iridium staining (Fluidigm Inc., 1:2000 concentration in 0.5 ml, RT). Finally, fixed samples were washed 3 times with DDW immediately prior to acquisition with CyTOF1 machine (Fluidigm Inc.).

### Trajectory assembly of healthy bone marrow

Manually gated Lin^-^HLADR^+^CD33^+^ cells of healthy individuals were pooled and filtered to include only those cells that express at least one myeloid marker from the following list: CD34, CD117, CD123, CD64, CD14 or CD11b. Of these cells, we randomly sampled 10,000 single cells on which we applied the diffusion-maps algorithm using the expression of the markers: CD38, CD34, CD33, CD64, CD14, CD13, CD117, CD11b and CD45. Pseudotime scores were calculated using the diffusion pseudotime algorithm^20^ (*destiny* R package). Pseudotime scores of the other non-sampled cells were calculated per cell as the average pseudotime scores of its 10-nearest neighbors relative to the markers set used for trajectory assembly. Resulting pseudotime scores of all the cells were scaled to [0,1] range. To avoid the density effects on interpolation of markers expression, we first subsampled a subset of cells randomly with inverse correlation to their density along the trajectory to achieve a uniform distribution of pseudotime scores on which interpolation was performed.

### Mapping of AML cells onto the healthy trajectory using devMap

Manually gated Lin-HLADR^+^CD33^+^ cells were filtered to include only those cells that express at least one myeloid marker from the following list: CD34, CD117, CD123, CD64, CD14 or CD11b. *Patient-specific trajectory assembly*. Per patient, 1,000 cells were first subsampled from the patient’s samples in an unbiased manner to achieve an equal representation of each one of the patient’s samples. To avoid the effects of the highly biased density in AML samples, ordering of the subsampled cells along a developmental trajectory was achieved by applying PCA with the following markers: CD38, CD34, CD33, CD64, CD14, CD13, CD117, CD11b and CD45 and using the first principal component axis as the pseudotime axis. Out of 17 patients, the PCA of 16 patients could be used for inference of a continuous meaningful trajectory. Those single cells from the patient’s samples that were not used for trajectory building were mapped to the assembled trajectory by calculating per cell the mean pseudotime values of its 10-nearest neighbors relative to the markers set used for trajectory assembly. Resulting pseudotime values of all cells were scaled to [0,1] range. To avoid the density effects on interpolation of markers expression, we first subsampled a subset of cells randomly with inverse correlation to their density along the trajectory to achieve a uniform distribution of pseudotime scores on which interpolation was performed. *Markers selection*. We defined the alignment quality as the mean cost per alignment step, calculated as the alignment-based distance divided by the alignment’s length (the number of steps taken for the alignment). For initial alignment, we chose a broad set of myeloid related markers whose expression vary along the trajectory: CD38, CD34, CD123, CD33, CD45RA, CD64, CD13, CD14, CD117, CD11b, CD45. We iteratively removed one marker at a time and re-aligned the AML and the healthy trajectories, each time assessing the alignment quality. The marker whose removal resulted with the greatest decrease in the mean cost of alignment step, was removed from the markers set. This process was repeated until the set of markers included only one marker. To identify the subset of markers that yield a high-quality alignment, at each step the alignment quality was compared to the one obtained from aligning each two healthy individuals with one another using the same set of markers. Assuming that the monocytic developmental trajectories are conserved across healthy individuals, we used the 90^th^ percentile of the mean cost per alignment step calculated between each two healthy individuals as a threshold. The largest set of markers under which the alignment of AML with the healthy trajectory resulted with an alignment quality smaller than the reference threshold was selected. *Global alignment*. Global alignment was applied using the *cellAlign* R package with the symmetric1 step pattern, without scaling of the dissimilarity matrix. *Alignment-based Mapping of AML single cells onto the healthy trajectory*. Single AML cells were assigned with the pseudotime values of the healthy interpolated point to which their closest interpolated point was aligned.

### Simulation for mapping performance demonstration

Synthetic AML data were generated by two methodologies. The first involved biased sampling of 5,000 healthy cells preferentially from different locations along the trajectory, based on the following probability distribution: *P_cell_* = *e*^−3∙|*pt*−*x*|^, where pt is the pseudotime of the cell and x is the location along the trajectory towards which the sampling is biased. Pseudotime scores were calculated as described before for the AML samples. Mapping accuracy per single cell was assessed by calculating the absolute differences between the original pseudotime values and those obtained under the biased sampling either before or after mapping by devMap. In the second methodology, synthetic datasets were generated on a per-marker basis by replacing the vector of the marker’s expression levels across cells with the one obtained by reversing the pseudotime based ordering. Pseudotime scores were calculated as described before for the AML samples, followed by application of devMap marker-selection step.

### Mapping of Levin et al dataset onto the trajectory

Raw fcs files were downloaded from https://community.cytobank.org/cytobank/. *Mapping of healthy individuals*. Manually gated Lin-HLADR^+^ cells were filtered to include only those cells that express at least one myeloid marker from the following list: CD34, CD117, CD64 or CD11b. A trajectory was generated for each individual as the first principal component axis in the PCA applied on 1,000 randomly sampled cells and using the markers: CD38, CD34, CD64, CD117, CD11b and CD45. Trajectory scaling, mapping of the other cells onto it, interpolation, alignment and maturation index calculation were performed as described above. *Mapping of AML patients*. All preprocessing steps are identical to the ones taken for the healthy individuals. Markers selection and single cell mapping were performed as described above. For identification of a high-quality alignment, we assumed that the monocytic developmental trajectories are conserved across healthy individuals, while taking into account the gap stemming from technical variability between the datasets. Specifically, we aligned every two healthy individuals, one from each dataset, and used the 90^th^ percentile of the mean cost per alignment step calculated across alignments as a threshold on alignment quality when aligning the AML sample to the healthy backbone.

---

**Supplementary Figure 1.**
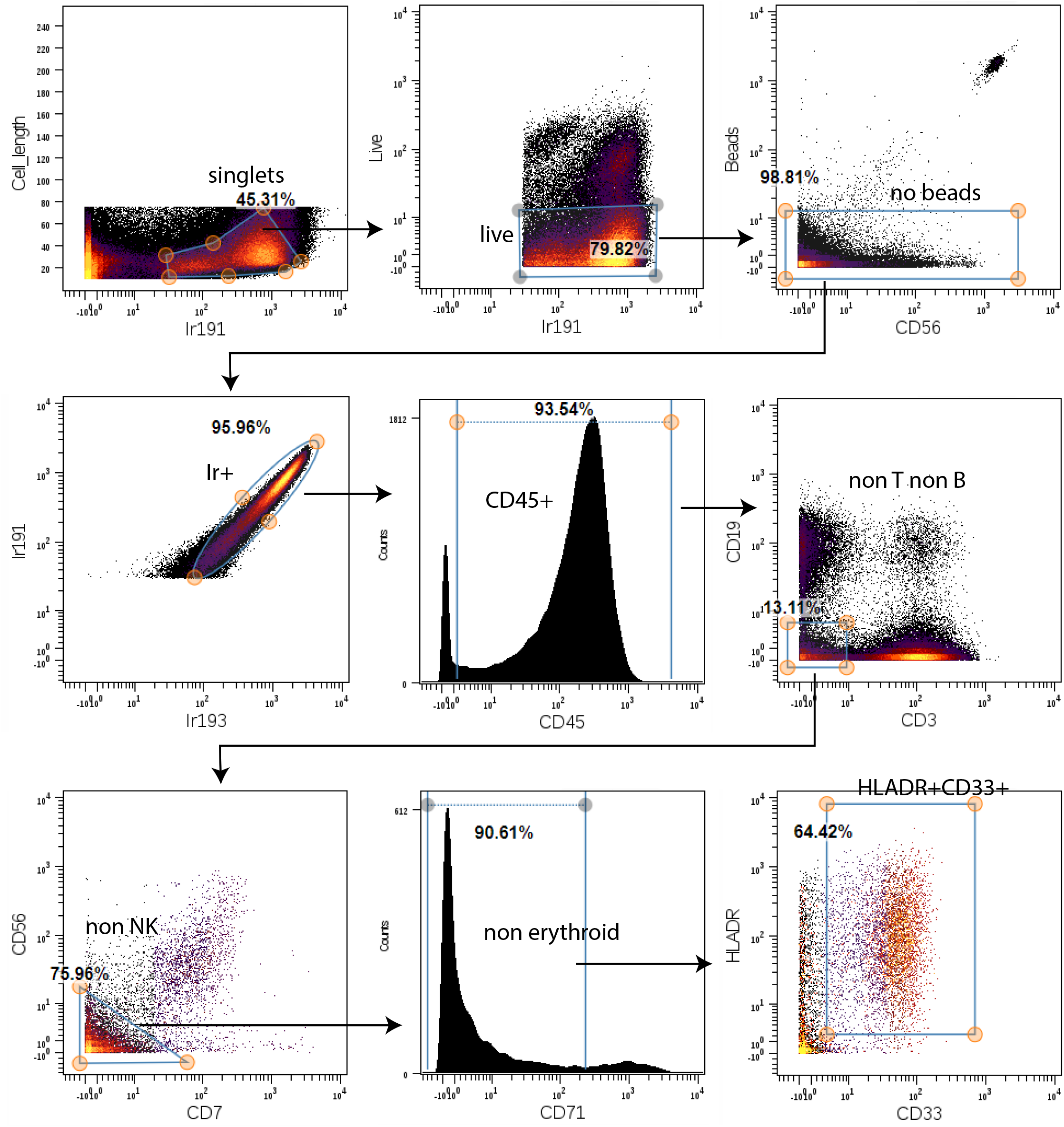
Gating scheme. Arrows link cell-populations to their parent population. Corresponding cell populations frequencies appear next to the gate.

**Supplementary Figure 2.**
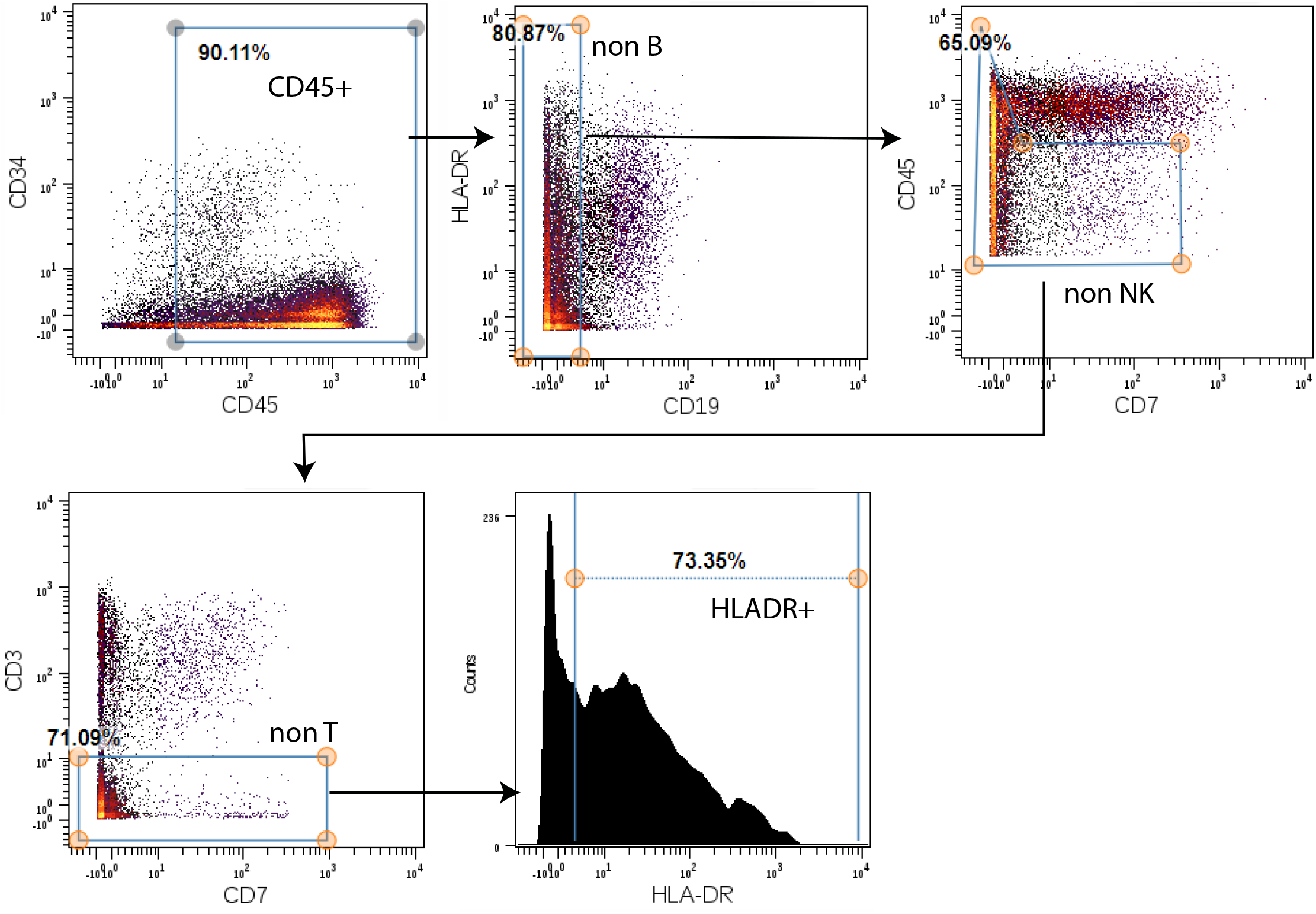
Gating scheme – Levin et al dataset. Arrows link cell-populations to their parent population. Corresponding cell populations frequencies appear next to the gate.

**Supplementary Figure 3.**
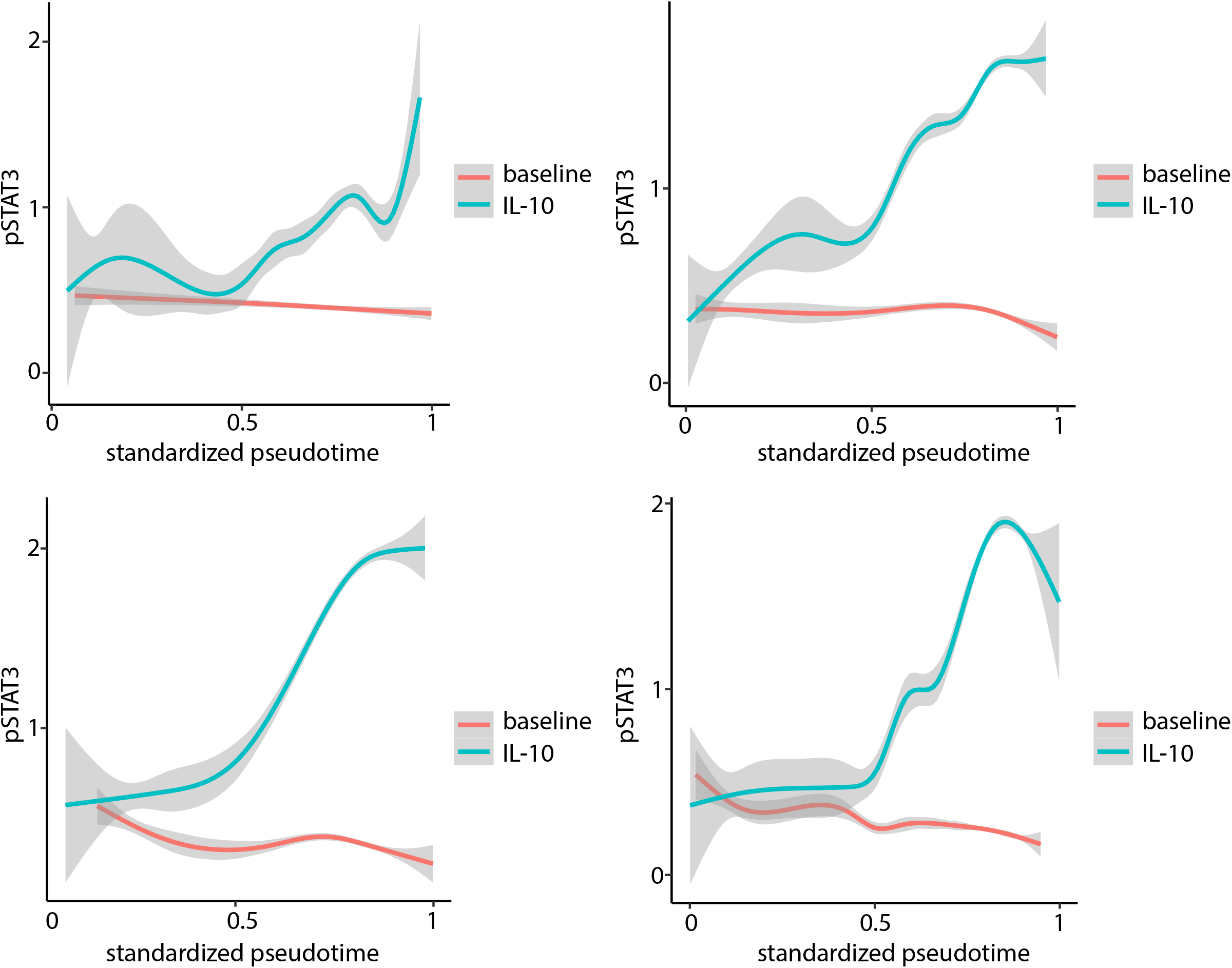
STAT3 phosphorylation smoothed expression levels as measured in healthy individuals from Levin et al dataset along standardized pseudo −10 stimulation (blue).

